# The suppressive potential of a gene drive in populations of invasive social wasps is currently limited

**DOI:** 10.1101/2022.06.27.497711

**Authors:** Adriaan B. Meiborg, Nicky R. Faber, Benjamin A. Taylor, Brock A. Harpur, Gregor Gorjanc

## Abstract

Social insects are very successful invasive species, and the continued increase of global trade and transportation has exacerbated this problem. The yellow-legged hornet, *Vespa velutina nigrithorax* (henceforth Asian hornet), is drastically expanding its range in Western Europe. As an apex insect predator, this hornet poses a serious threat to the honey bee industry and endemic pollinators. Current suppression methods have proven too inefficient and expensive to limit its spread. Gene drives might be an effective tool to control this species, but their use has not yet been thoroughly investigated in social insects. Here, we built a model that matches the hornet’s life history and modelled the effect of different gene drive scenarios on an established invasive population. To test the broader applicability and sensitivity of the model, we also incorporated the invasive European paper wasp *Polistes dominula*. We find that although a gene drive can spread through a social wasp population, it can only do so under stringent gene drive-specific conditions. The main issue is that the large number of offspring that social wasp colonies produce guarantees that, even with very limited formation of resistance alleles, such alleles will quickly spread and rescue the population. Furthermore, we find that only a gene drive targeting female fertility is promising for population control due to the haplodiploidy of social insects. Nevertheless, continued improvements in gene drive technology may make it a promising method for the control of invasive social insects.

## Introduction

Invasive species represent a global issue that has worsened with increased global trade and transportation (1, 2). Suppression of these invasive species is often prohibitively expensive, labour intensive, and largely ineffective (1). One such species currently invasive in Europe is the yellow-legged hornet *Vespa velutina nigrithorax*, hereafter called the Asian hornet. This insect was introduced to France from Southern China in 2004 (3, 4) and quickly spread to the whole of France, most of the Atlantic coast, Northern Italy, Belgium, parts of the United Kingdom, and parts of Germany, where the northernmost finding was made in 2020 (5, 6). This spread is in accordance with previous modelling of suitable environments (7, 8). Modelling also showed that there are many more areas in Europe suitable for the Asian hornet to invade (8).

The Asian hornet likely has a serious impact on commercial bee colonies (9–11) and potentially on other pollinators such as wild bees and syrphids (5, 11). Up to two thirds of its diet consists of honey bees (11), and the annual loss for France’s honey and pollination industry in 2015 was estimated at 53.3 million euros (8). The invasion probably started with only a single fertilised queen (3), which underscores the great invasive potential of the Asian hornet. Indeed, one queen produces on average 300 gynes that all have the potential to start a new nest the next year (12). On top of this great reproductive potential, controlling the Asian hornet through conventional means is difficult; the nests are high up in trees, and thus hard to find, and they are also hazardous to approach. Bait trapping with food or chemicals is currently the most reliable control method, though it is only partially effective and is not species-specific (13). These inadequacies highlight the need to find an effective strategy to control the Asian hornet before it spreads to other suitable regions.

Over the last decade, gene drives have emerged as a potential tool to control invasive populations for which other measures are ineffective (14–18). A gene drive is a genetic element that spreads through a population over generations at a super-Mendelian rate. For population control, it is engineered to impose a fitness cost once it is prevalent. For example, a gene drive may disrupt a haplo-sufficient female fertility gene. Haplo-sufficient means that a single functioning copy of the gene is enough for a female to be fully fertile. At first, the gene drive is present mostly in a heterozygous state due to matings with wildtype individuals. Once it reaches a higher frequency, matings between gene drive individuals will occur more frequently and offspring will be homozygous for the gene drive, with female offspring thus being infertile. This way, population fecundity declines through the reduced fertility of homozygous individuals (15). Gene drives have been demonstrated to work in yeast (19), fruit flies (20), mosquitoes (21, 22), and mice to a lesser extent (23, 24). The field has recently focused on improving safety and containment which would make gene drives controlled enough for release in the wild (25). A range of gene drives have been designed to be less invasive by nature, or to mitigate risks to an extent. Some stop spreading after a certain number of generations and are thus self-limiting (26–28), some require high introduction frequencies (29–31), some can stop or remove a gene drive that is already present in a population (32, 33), and some target locally fixed alleles so that the gene drive cannot spread in non-target populations (34). With the advance of such containable gene drives, we can start to consider gene drive technology as a potential tool for controlling invasive social insects like the Asian hornet.

Creating a realistic life history model for a population is one of the first steps in determining if a gene drive might be an effective control agent (35). A previous study has modelled a gene drive causing male sterility in another haplodiploid social hymenopteran, the common wasp *Vespula vulgaris*, that is invasive in New Zealand (36). This type of gene drive was shown to be only mildly promising, because there was a trade-off between the spread and the impact of the gene drive. Namely, if the gene drive causes complete male sterility, it is unlikely to spread, whereas if the gene drive causes incomplete male sterility, it is unable to impact the fertility sufficiently for population control (36). Therefore, this specific gene drive design works similarly to sterile insect technology: a powerful, but only temporary method. Therefore, other gene drive designs are needed to control such populations more efficiently (37). For example, three recent modelling studies showed that gene drives can work in haplodiploid species under specific conditions (38–40).

In this study, we model several gene drive strategies to investigate the potential of gene drive technology to control Asian hornet populations. We look into both gene drive parameters and life history traits to find the most important factors that could affect the success of gene drives to control these invasive populations. We developed a model that can be easily adapted to other social haplodiploid insects, as the Asian hornet is not the only invasive social hymenopteran. Indeed, this group of insects, which is comprised of wasps, bees, and ants, contains many successful invasive species (41). We demonstrate the flexibility of the model by modelling a second invasive social hymenopteran, *Polistes dominula*, hereafter called the European paper wasp. This paper wasp is a widespread invasive species with a very different biology than the Asian hornet. It has much smaller colonies, and queens are almost exclusively monogamous (42). Modelling two species across the social wasp spectrum showed how life history influences gene drive efficiency. Our results show that gene drives can be used to contain an invasive population of either species, but only when the gene drive achieves extremely high efficiency. This is the case because high per-colony offspring numbers allow every possible resistance allele to subsist in the population once they arise, limiting the success of gene drives that generate resistance alleles. Consequently, species with lower numbers of offspring require a slightly less stringent gene drive efficiency. Therefore, we need significant advances in gene drive technology, especially lower rates of non-homologous end-joining and more efficient cutting rates, before this technology can control populations of invasive social wasps.

## Material and Methods

The model is an individual-based, stochastic, year-by-year model with two main parts: 1) a realistic model of a social wasp population based on their life history and 2) the implementation of a gene drive (see figure 1). The model was built using R version 4.1.2 (43). We use AlphaSimR package version 1.0.1 (44). This package is designed for animal and plant breeding studies, but also serves as an excellent tool for modeling population genetics studies. The model can be found on GitHub at https://github.com/HighlanderLab/ymeiborg_hornet_gd.

**Fig. 1.**
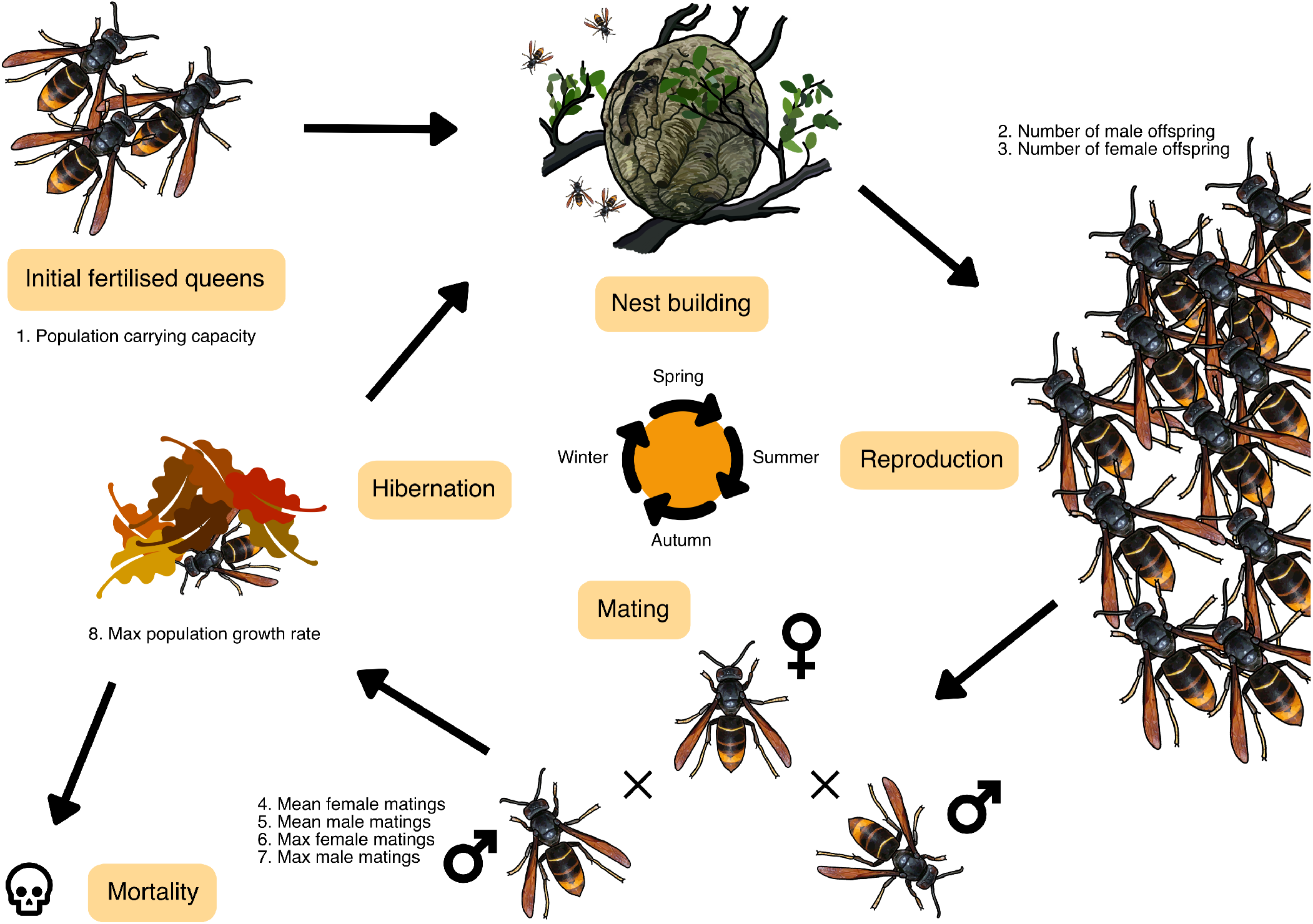
Overview of the demographic model. For a specific explanation of the model, see the Material and Methods.

### A. Model structure

The structure of our model mimics the life cycle of a typical haplodiploid social wasp. Throughout the model, we only track female numbers and genotypes, as males only live for a short amount of time for mating purposes and little is known about their survival. We initiate the model with an equal number of females and males, of which the females (queens) represent the starting population size. Since we initiate the model with mated queens, each male is assigned as a mate for a queen. In the scenario of releasing females with gene drive, these are added as well. In the scenario of releasing males with a gene drive, these are added in the mating step in the first generation of the model. Depending on the chosen number of generations to model, the following steps are repeated:

1. **Offspring generation**. First, offspring are generated for each successfully mated queen. This is the point at which gene drive dynamics, such as homing or resistance allele formation, take place in the queen’s germline as described in section B. Haploid male offspring are produced by the queen from unfertilised eggs, while diploid female offspring originate from fertilised eggs. For each female that has mated more than once, the total number of offspring is divided randomly over the multiple mates. We assume an equal number of female and male offspring. Previous findings report a 3:1 male to female ratio in invasive populations (45), but this is likely a result of overproducing infertile diploid males under inbreeding (46), as non-Fisherian sex ratios are unlikely to persist at a population level (47, 48). The number of female and male offspring are used as averages for a Poisson distribution for each sex so there is natural variation in the number of female and male offspring from each queen. For Asian hornets, on average, one queen produces around 300 females (12) and we assume the same number for males. European paper wasp nests are much less productive, with a single nest producing ~20 offspring of each sex (49).
2. **Mating**. We have implemented random mating between males and females, except we impose the fitness costs of having a gene drive here. Both males and females with a gene drive in their genome have a certain probability of being removed from the mating pool. We assumed full dominance for the fitness cost, so both heterozygous or homozygous individuals are equally affected. Next, as polyandry is shared by many social hymenoptera (50, 51), we have included it in the model. For both females and males, the model takes the mean number of matings, the minimum number of matings, the maximum number of matings, and based on these values generates a random number of matings from a truncated Poisson distribution. For males, the mating frequencies from (52) include a high frequency of nonmating males, so we have not zero-truncated their mating rate distribution. The Asian hornet queen is polyandrous and is able to attract mates at great distances (53), so for mating rate we use a zero- and max-truncated Poisson distribution with a mean of 3.275 and a maximum of 4 (3). Male Asian hornet mating rates are unknown, so we used a max-truncated Poisson distribution with a mean of 0.9 and a maximum of 2 based on mating frequencies for *Vespula maculifrons* (52). Mating rates for European paper wasp females are estimated to fall between 1 and 1.1 (54, 55), and as such were modeled using a zero- and max-truncated Poisson distribution with a mean of 0.2 and a maximum of 2. There is no accurate data that capture total male reproductive output in *P. dominula* either, so we used the rates reported in *V. malicifrons* (52) instead. After mates have been randomly selected this way, we remove pairings of which one or both individuals are infertile due to the gene drive. We thus assume that neither female or male knows whether their mate is fertile until the offspring generation, at which point it is too late to find a new mate. This assumption is in line with the observation that hymenopteran females are largely unable to discriminate against infertile but otherwise healthy diploid males (46).
3. **Mortality**. Population size is regulated using a Poisson distribution around a logistic function, which ensures a maximum population growth rate and carrying capacity (56):

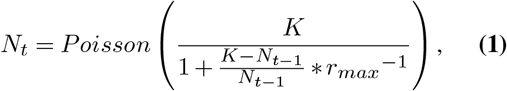

where *N_t_* is the number of females in generation *t, N*_*t*–1_ the number of females in generation *t* – 1, *K* is the carrying capacity, and *r_max_* is the maximum growth rate. Note that the maximum growth rate *r_max_* is calculated using 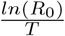 with *R*_0_ being the intrinsic growth rate observed by (56), *T* being the generation time of 1 year. Then, using the maximum possible population size in this generation and total number of female offspring, we calculate the mortality rate:

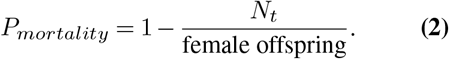 Each female thus has a probability of dying. The surviving females become queens and generate the offspring for the next generation. Male population numbers are not monitored, as they only live for a short amount of time and little is known about their survival.

### B. Gene drive implementation

Although AlphaSimR was designed to model large numbers of loci for quantitative genetics in plant and animal breeding, the framework is perfect for tracking the single locus of a gene drive. Each individual is modelled with a single gene drive locus and inheritance is random following Mendelian laws. We have implemented a basic homing gene drive which copies itself to the other chromosome in the germ line and has four potential alleles: wildtype (WT), gene drive (GD), resistance (RE), and non-functional (NF). We model diploid females and haploid males, so there is no gene drive activity in the male genome. Like Prowse et al. (2017) (37), we account for a probability of cutting (*P_cut_*) of 0.95, a probability of non-homologous end-joining (*P_NHEJ_*), and a probability that nonfunctional repair occurs (*P_NFR_*) of 0.67, which is the probability of a frame-shift occurring within the targeted gene. We evaluate the sensitivity of results to the default probability of cutting (*P_cut_*) and probability of non-homologous end-joining (*P_NHEJ_*). We also model a fitness cost of carrying the gene drive abstracted as a certain probability of mortality before mating (*P_mort_*). We conservatively assume that the fitness cost of carrying the gene drive has complete dominance, that is, it is equally deleterious for homo-, hemi-, and heterozygotes.

### C. Modelled scenarios

To show two sides of the social wasp spectrum, we do all our modelling for two species of invasive social wasps that have distinct life histories. The first species we model is the Asian hornet, which is an extremely successful invasive species, probably due to its ability to produce many new queens each generation. Females of this species are also very polyandrous. The second species is the European paper wasp, which has a more modest number of offspring and females are less polyandrous.

We first focus on which type of homing gene drive might work best for social wasps. We test a neutral gene drive and gene drives that cause female, male, and both-sex infertility. We test all of these gene drives with or without polyandry in both species. In these scenarios, we model current standard gene drive efficiencies: *P_cut_* = 0.95 and *P_NHEJ_* = 0.02, keeping *P_mort_* at 0, as in (37).

Next, we vary the parameters of the gene drive to find the space in which a gene drive works in our species. First, we vary *P_cut_* and *P_NHEJ_* together, second we vary *P_mort_* and *P_NHEJ_* together, and third we vary *P_cut_* and *P_mort_* together. Finally, we vary life history parameters of the wasps using “optimal”, “intermediate”, and “realistic” gene drive parameters, to evaluate how these parameters influence gene drive performance. The optimal gene drive scenario uses *P_cut_* = 1, *P_NHEJ_* = 0, and *P_mort_* = 0. The intermediate gene drive scenario uses *P_cut_* = 0.97, *P_NHEJ_* = 0, and *P_mort_* = 0.1. The realistic gene drive scenario uses *P_cut_* = 0.95, *P_NHEJ_* = 0.02, and *P_mort_* = 0.1.

For each model, except figure 2 and S2, we ran 10 replicates to get an estimate of the variance of the results. For figures 2 and S2, we ran 100 repetitions because the male gene drive release showed a lot of variation.

**Fig. 2.**
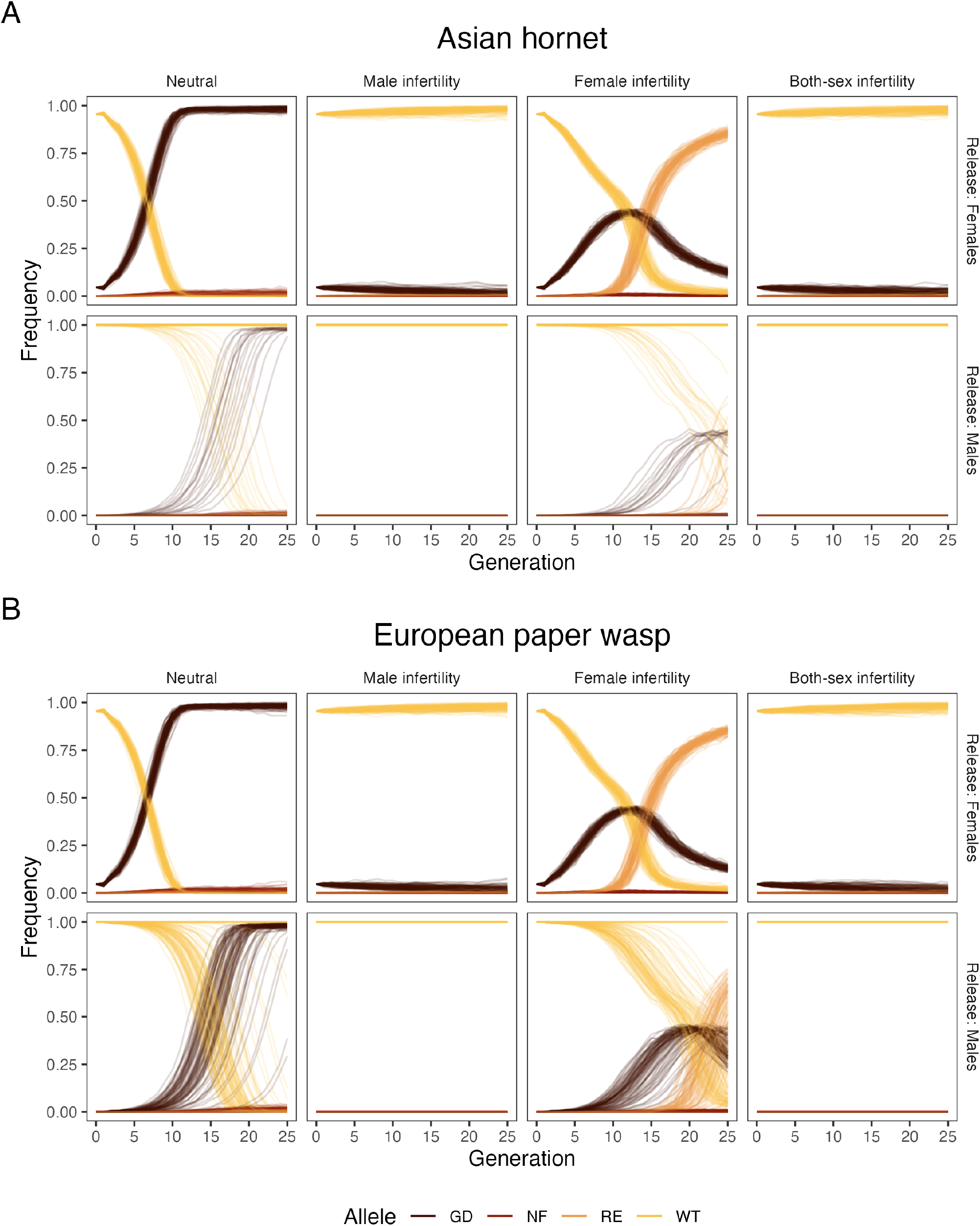
Frequencies of wildtype (WT), gene drive (GD), resistance (RE), and non-functional (NF) alleles in a female Asian hornet population (**A**) and a female European paper wasp population (**B**) by gene drive strategy and release carriers (females or males). The different strategies (neutral, male infertility, female infertility, and both-sex infertility) determine how the gene drive operates. In the neutral strategy there is no fitness cost to having the gene drive, whereas in the infertility strategies, the designated sex cannot reproduce when homo- or hemizygous for the gene drive.

## Results

The spread of the gene drive depends heavily on its design. In the following paragraph we describe the frequencies of wild-type and gene drive alleles shown in figure 2, while population sizes for the same scenarios are available in figure S1. A neutral gene drive spreads quickly through the female Asian hornet population, but does not reach complete fixation as resistance alleles and nonfunctional alleles appear (figure 2A). These resistance alleles and non-functional alleles are then subject to random drift. Female carriers are much more effective in spreading the gene drive (figure 2A-top row) than male carriers (figure 2A-bottom row). More often than not, the gene drive introduced by males goes extinct (in 83 out of 100 replicates), particularly when coupled with male infertility (extinction in all 100 replicates). In the cases it does spread, the spread is slower and less consistent compared to an introduction through females.

Gene drives designed for population suppression show similar dynamics. When the gene drive targets fertility, we see that targeting male fertility prevented spread of the gene drive through the population. The spread stops immediately when the gene drive is introduced via males. This effect persists when targeting female fertility at the same time. A gene drive targeting female fertility spreads rapidly through the population, but also leads to a rapid increase of resistance alleles in the population, which overtake the gene drive.

The spread of the gene drives does not differ much between the European paper wasp (figure 2B) and the Asian hornet (figure 2A). As in the Asian hornet, targeting male fertility prevents the gene drive from spreading, including when coupled to female infertility. Targeting female fertility remains more effective. European paper wasp males are more successful at introducing a gene drive (11 out of 100 replicates failed to spread the gene drive) than the Asian hornet males, but females remain more effective nonetheless.

To untangle the effects of polyandry and reproduction rate on the spread of the gene drive, we modeled the Asian hornet without polyandry, and the European paper wasp with polyandry (figure S2). In the latter case the female can mate with up to 4 males. Comparing figures 2 and S2 the effect can mostly be seen when males are used to release the gene drive. The top rows (female release) of figure 2A and B and figure S2A and B do not differ in any discernible way. The bottom rows (male release) of figure 2A and B and figure S2A and B are different, for the neutral and female infertility gene drive. When polyandry is introduced in the European paper wasp (figure S3B), the gene drive spreads more reliably when released by male carriers. Conversely, the variability in the spread of the gene drive increases when polyandry is removed from the Asian hornet (figure S2A). However, although gene drive spread via male carriers becomes more reliable when species are polyandrous, release via female carriers clearly remains the more reliable option.

Non-neutral gene drives are not capable of fixing in any of the modelled populations (figure 2 and S2). This is because resistance alleles rapidly spread through the population due to the heavy selective pressure the gene drive introduces. This way, these resistance alleles outcompete the gene drive, and rescue the population. This is so much the case that non-neutral gene drive do not change the population sizes (figure S1). Resistance alleles are formed when the DNA strand is repaired with non-homologous end-joining after being cut by the Cas9 protein. This method is error-prone and can insert viable mutations which decrease the affinity of the gRNA for the target sequence. Probability of non-homologous endjoining (*P_NHEJ_*) is not the only parameter inherent to the gene drive that affects its efficiency. Different gRNA target sites can have different probabilities of cutting the opposite DNA (*P_cut_*). Gene drives have an inherent fitness cost when present in a genome, here called the probability of heterozygous mortality (*P_mort_*).

We varied gene drive parameters and estimated which parameters are critical and what values are required for a proper population suppression. In both the Asian hornet and the European paper wasp, only absence of non-homologous endjoining (*P_NHEJ_ =* 0) will make gene drive work reliably (figure 3A-D). Nor will a cutting rate (*P_cut_*) below 1 make gene drive work reliably in the Asian hornet (figure 3A). In the European paper wasp, a cutting rate of 0.97 is the lowest workable value (figure 3B). Heterozygous mortality (*P_mort_*) affects both species nearly equally. Reliable suppression stops above a 0.275 (figure 3C and D) in both cases, although the drop in reliability is steeper in the Asian hornet. We also explored cutting rate against heterozygote mortality in a gene drive in the absence of non-homologous end-joining (figure 3E and 3F). The European paper wasp has a larger area of success than the Asian hornet. Estimates for the European paper wasp have more noise, which could be due to lower reproductive values. Based on these results we have chosen three different gene drive conditions for further analysis: optimal, intermediate, and realistic as shown on figure 3.

**Fig. 3.**
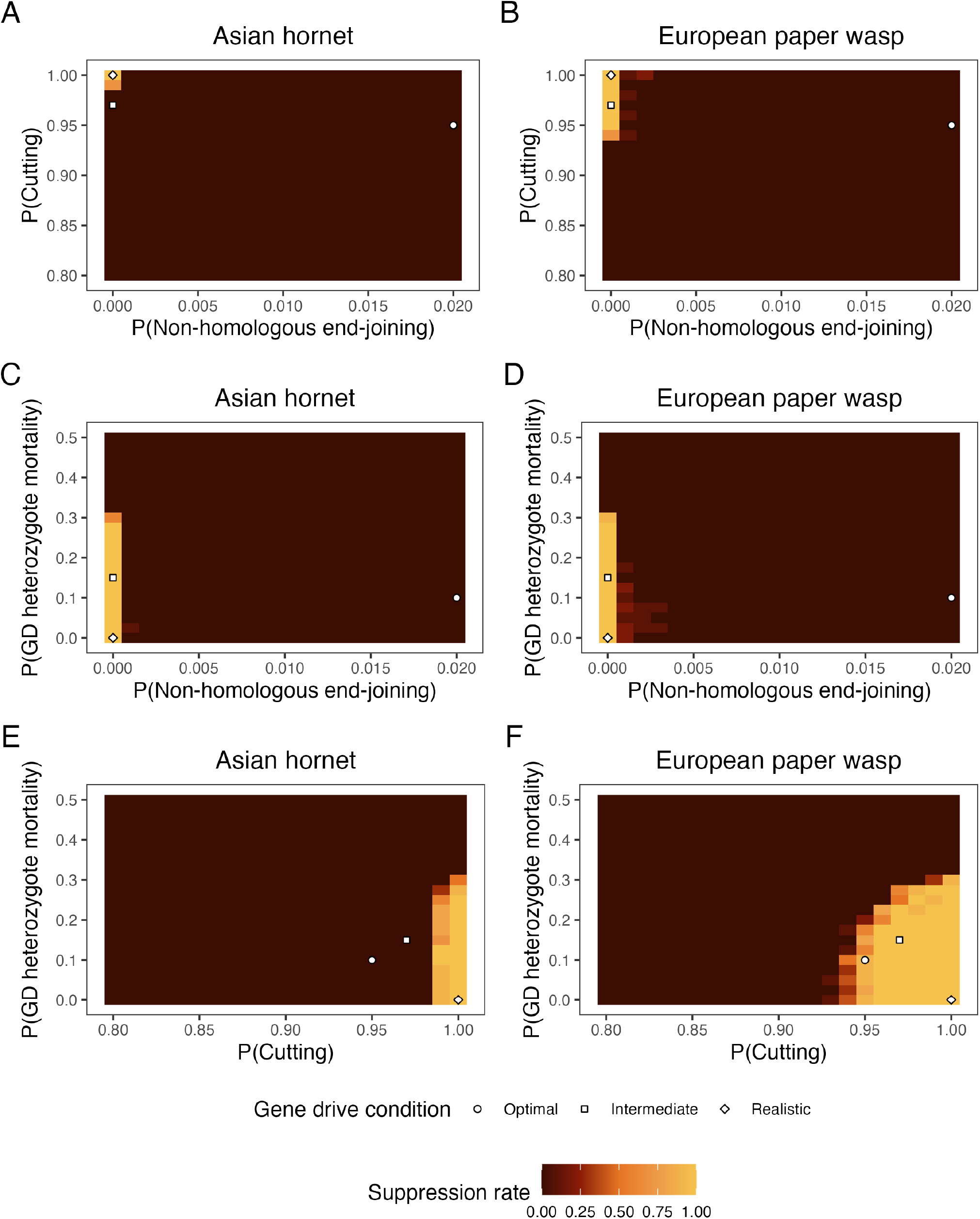
Heatmaps of the suppression rate in the Asian hornet (**A** & **C**) and the European paper wasp (**B** & **D**) using gene drives that have varying probabilities of nonhomologous end-joining (P(Non-homologous end-joining) or *P_NHEJ_*) and cutting (P(Cutting) or *P_cut_*) or mortality of gene drive heterozygotes (P(GD heterozygote mortality) or *P_mort_*). Note that the model was run for 50 generations because in many intermediate cases 25 generations was not sufficient for the gene drive to suppress the population.

To test the sensitivity of our model and results to the used biological parameters, we performed a sensitivity analysis for both modeled species (figure S3 and S4) with the three different gene drive conditions: optimal, intermediate, and realistic. Under realistic conditions, there is only a visible effect in the European paper regarding the number of offspring parameter. Only an extremely low mean number of progeny affects the spread of the gene drive (figure S4). The other parameters have no effect on the gene drive success in either species, which is in line with figure 2. Resistance alleles prevent the fixation of the gene drive and rescue the population. The intermediate condition was between the optimal and realistic conditions and showed that in the European paper wasp all the tested parameters influence gene drive success, particularly the mean number of progeny. In the Asian hornet, there is only a small effect visible for the mean number of progeny. The gene drive is unable to suppress the population in any of the other scenarios. Since this parameter differs the most between the two species and it is critical for the spread of alleles under drift, we evaluated a wide range of values in each species, well beyond the biological limits in the two species (figure S5). This sensitivity analysis showed a similar effect of the number of progeny on gene drive success in both species irrespective of their other biological parameter values (figure S3 and S4). This indicates that under the intermediate gene drive conditions the sheer number of progeny in Asian hornets is a limiting factor for the capacity of gene drives to suppress this invasive species. Under optimal conditions, the gene drive is always successful, regardless of the biological parameters.

## Discussion

Invasive social hymenopterans, such as the Asian hornet, can cause significant ecological and commercial damage in their invasive ranges. Difficulties in the application of conventional methods for the control of invasive social hymenopterans have led to a surge of interest in the possibility of using gene drive to suppress such invasive populations. In this paper we have modelled the efficacy of gene drives for population control of two invasive social wasps, the Asian hornet (*Vespa velutina nigrithorax*) and the European paper wasp (*Polistes dominula*) under a variety of gene drive and life history parameters. We find that the fecundity of social insect colonies represents a major limiting factor for the efficacy of gene drives. As a result, gene drives can only suppress invasive wasp populations when gene drives operate with extreme efficiency. Other life history traits that we examined were less impactful. We also find that a gene drive targeting fertility is only effective when targeting females. When a gene drive targeted male fertility or the fertility of both sexes, it could not spread. Overall, our results indicate that, despite the potential value of gene drive as a control agent for invasive social insects, the current efficiency of gene drives is insufficient to overcome the biological efficiency of social insect reproduction.

We found that selection resulted in the rapid fixation of alleles conferring resistance to drive, such that the gene drive was never able to reach fixation under currently-realistic parameters of gene drive efficiency. While there are proposals that improve gene drive efficiency (for example 57), the relatively high fecundity of female social insects raises the bar for required gene drive efficiency. Due to the reversal of longevity/fecundity trade-offs, reproductive females in social hymenopteran species may reach levels of fecundity magnitudes higher than those possible for non-social organisms (58, 59). This fecundity reduces the effects of drift and therefore the potential efficacy of gene drive: the more progeny each gene drive carrier female produces, the more likely that all possible allele combinations will be represented among those progeny, thereby reducing the likelihood that any given allele will be fixed by chance. For the same reason, the probability that a novel resistance allele will be lost by drift before being spread by positive selection is reduced when female fecundity is higher.

Our comparison of two different social wasps with very different numbers of progeny demonstrates the importance of fecundity. The parameter space in which the gene drive was able to effectively suppress a population was larger for the European paper wasp than for the Asian hornet. This difference appears to have been driven by the lower mean progeny sizes in the paper wasp. Notably, when we varied mean progeny number for each species while keeping all other variables equal, the rate of suppression was approximately equal for the two species for any given number of progeny, suggesting that it is fecundity rather than other factors that drove the difference in gene drive suppressing the population between the species.

Our results indicate that, in social wasps, a driving element that targets fertility can only successfully spread if its effects are female-specific. Gene drives targeting fertility either in males or in both sexes were never successful in our modelling, because haploid males are unable to act as asymptomatic carriers of the gene drive. Our results therefore indicate that any successful attempt to control invasive haplodiploid species using fitness-targeting gene drives will necessarily target females rather than males, in line with the findings of a previous modeling study (39). This fact may represent a significant impediment to the biological control of invasive haplodiploids using gene drives, since the rate of formation of resistance alleles is expected to be substantially higher in females than in males (60).

High fecundity and haplodiploidy represent two significant challenges to the potential efficacy of gene drive as a control agent for invasive social wasps. A third challenge is the risk that the gene drive could spread to other wasp species by hybridization: rates of inter-specific hybridization appear to be significantly higher among hymenopterans than other arthropods (61), and the introgression of a driving allele into native wasp species is a meaningful risk given that social wasps perform important ecosystem services in their native ranges (62). Thus, even if the technical efficiency of drive can be optimised to the point that it is a viable option for the control of invasive wasps, gene drive’s safety as a control agent may remain a significant concern.

We found that a neutral gene drive could spread much more reliably than one targeting fertility. In theory, a gene drive without any directly detrimental phenotypic effects can still contribute to biocontrol, if the gene drive leaves affected individuals vulnerable to further control measures. For example, a gene drive that disrupts resistance to a specific pesticide could allow that pesticide to be used as a control measure once the gene drive has reached high frequency or become fixed (40). However, such an approach requires targeted management techniques, which are not currently available for invasive social hymenopterans excepting a few ant species (63, 64). As such, direct targeting of female fertility is likely to remain the most promising route for managing invasive social hymenopterans using gene drive.

Despite the challenges presented by haplodiploidy and high fecundity, we found that a gene drive targeting female fertility could spread to a significant frequency (~0.75) within social wasp populations before being negated by the spread of resistance alleles. Even at these high frequencies, however, we found no impact of gene drive on population size, because the high fecundity of individual reproductive females allowed the population to remain at carrying capacity even while the driving allele was present at high frequencies. Anything short of complete fixation of the gene drive was insufficient to suppress an invasive population. Combined with the rapid generation of resistance alleles in a short number of generations, this result indicates that a homing gene drive would be ineffective at suppressing invasive social wasp populations over both long and short timescales.

Like any model, ours includes assumptions and simplifications. Parameterisation of the model proved difficult due to a relative paucity of life history data for vespid wasps, despite the ecological importance of this group (62). For example, we assume that female mating rates are zero-truncated, such that females never fail to find at least one male with whom to mate. This assumption may become unrealistic for very small population sizes, but empirical data that would allow us to accurately model this effect are lacking. For this reason, we instead used estimates from several closely related species as described in our methods. Other potential limitations of our model include the lack of any spatial component, including a complete lack of immigration and emigration, the assumption of perfect admixture without mate choice, and the high heterozygous mortality.

## Conclusions

We have modeled the spread of a homing gene drive under a variety of conditions of life history and drive efficacy through populations of two invasive social wasps: the Asian hornet and the European paper wasp. We find that, due to large progeny numbers produced by reproductive females in these species, a homing allele can only reach fixation under extremely efficient drive conditions. These findings, together with limitations imposed by haplodiploidy and potential for inter-specific hybridization, highlight the difficulty of applying genetic biocontrol measures to social hymenopterans. We conclude that until it is possible to develop gene drives with much higher efficiency of spread and much lower rates of resistance allele formation, more conventional approaches such as nest destruction and bait trapping will remain the best methods for the control of invasive social wasps.

## ACKNOWLEDGEMENTS

This study was conceived of by all authors. Analyses were performed by ABM and NF. The manuscript was written by all authors.

ABM acknowledges funding from the European Molecular Biology Laboratory. NF acknowledges funding from the Graduate School for Production Ecology & Resource Conservation call 2020. BAT acknowledges funding from the USDA APHIS Farm Bill and a Human Frontiers in Science Program Postdoctoral fellowship. BAH acknowledges funding from the USDA APHIS Farm Bill. GG acknowledges the BBSRC Institute Strategic Programme funding to The Roslin Institute (BBS/E/D/30002275). For the purpose of open access, the authors have applied a CC BY public copyright licence to any Author Accepted Manuscript version arising from this submission.

## Supplementary Material

**Fig. S1.**
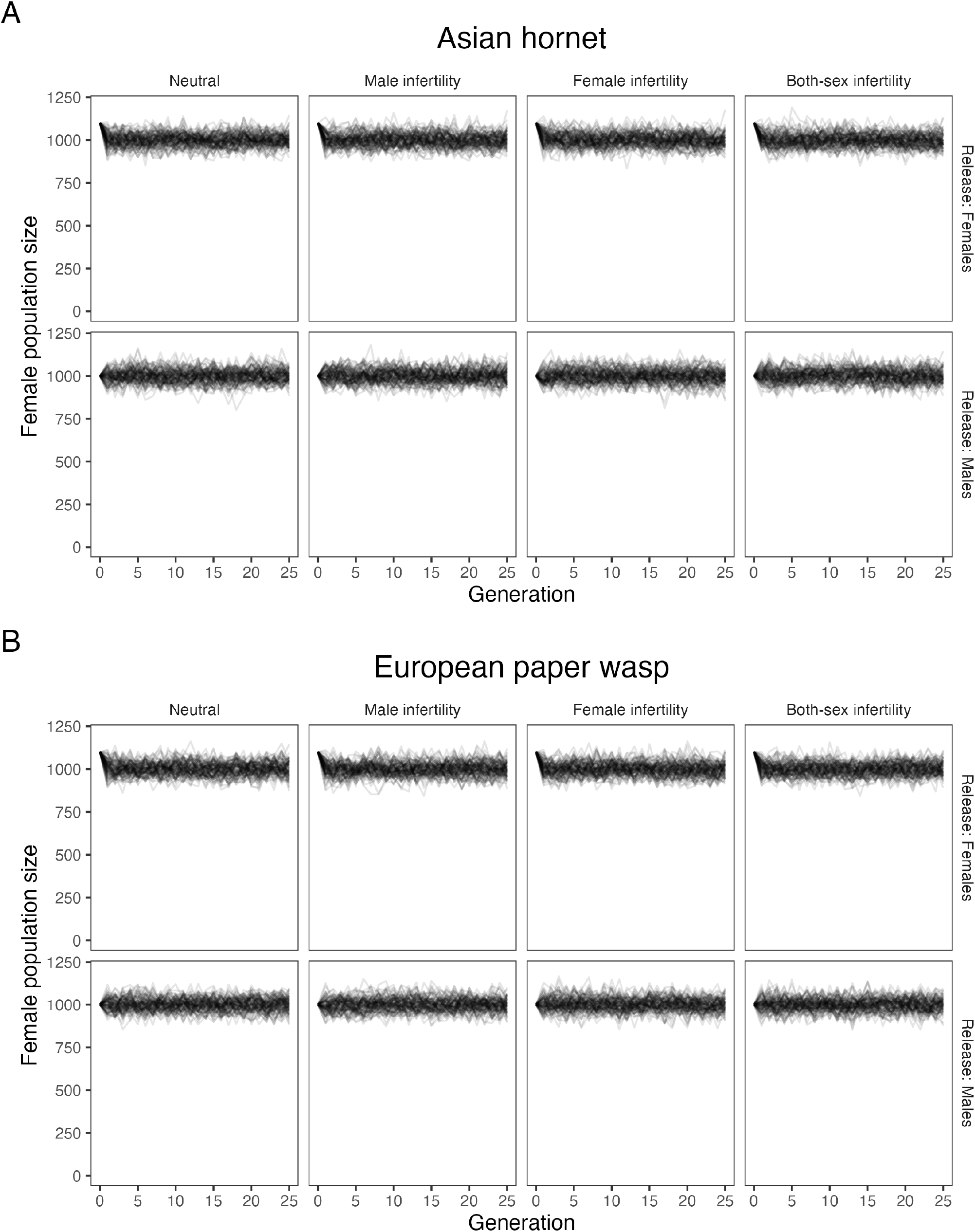
Female population size for Asian hornet (**A**) and European paper wasp (**B**) by gene drive strategy and release carriers (females or males). The different strategies (neutral, male infertility, female infertility, and both-sex infertility) determine how the gene drive operates. In the neutral strategy there is no fitness cost to having the gene drive, whereas in the infertility strategies, the designated sex cannot reproduce when homo- or hemizygous for the gene drive.

**Fig. S2.**
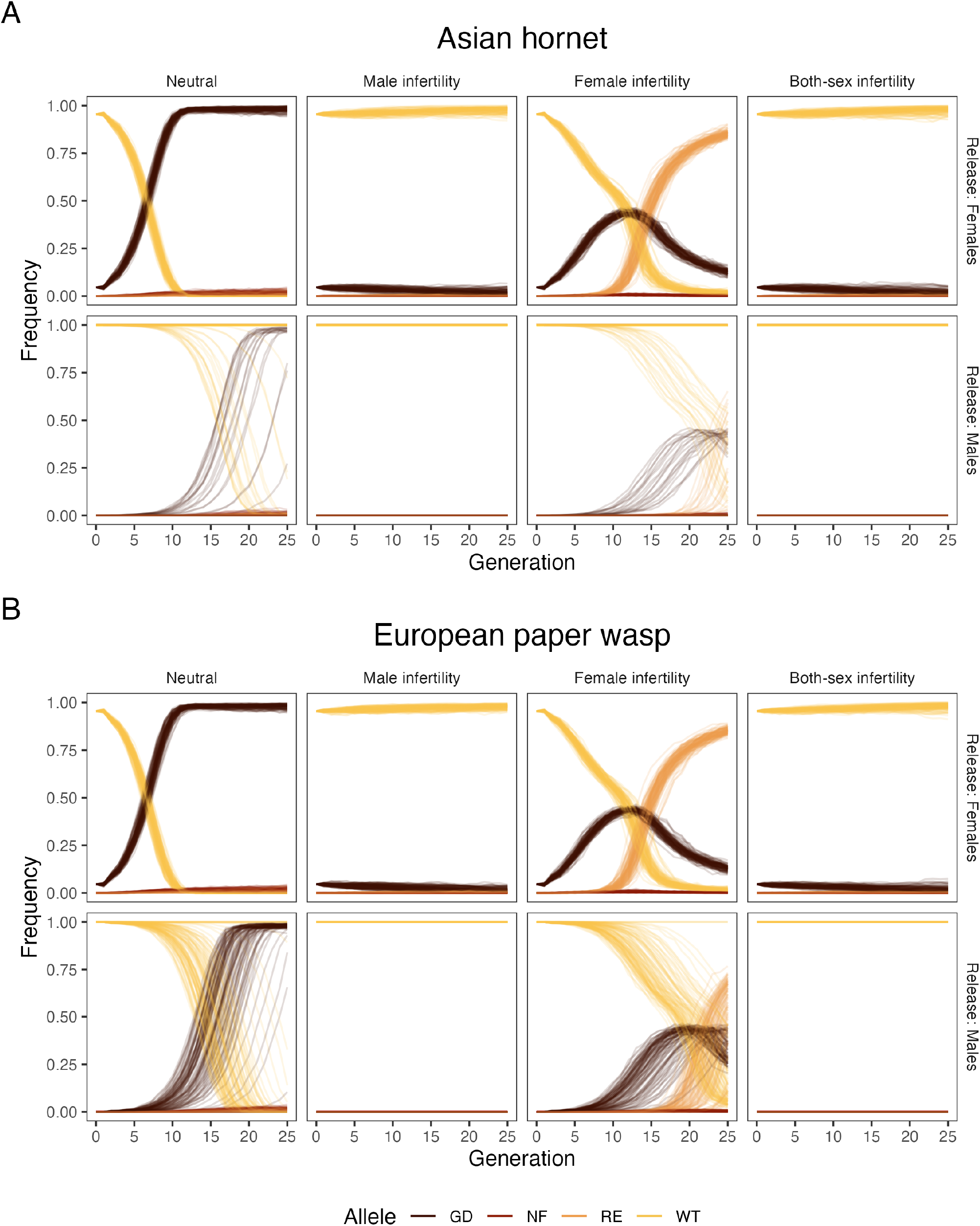
Frequencies of wildtype (WT), gene drive (GD), resistance (RE), and non-functional (NF) alleles in a female Asian hornet population with no polyandry (**A**), and a female European paper wasp population with polyandry (**B**) by gene drive strategy and release carriers (females or males). The different strategies (neutral, male infertility, female infertility, and both-sex infertility) determine how the gene drive operates. In the neutral strategy there is no fitness cost to having the gene drive, whereas in the infertility strategies, the designated sex cannot reproduce when homo- or hemizygous for the gene drive.

**Fig. S3.**
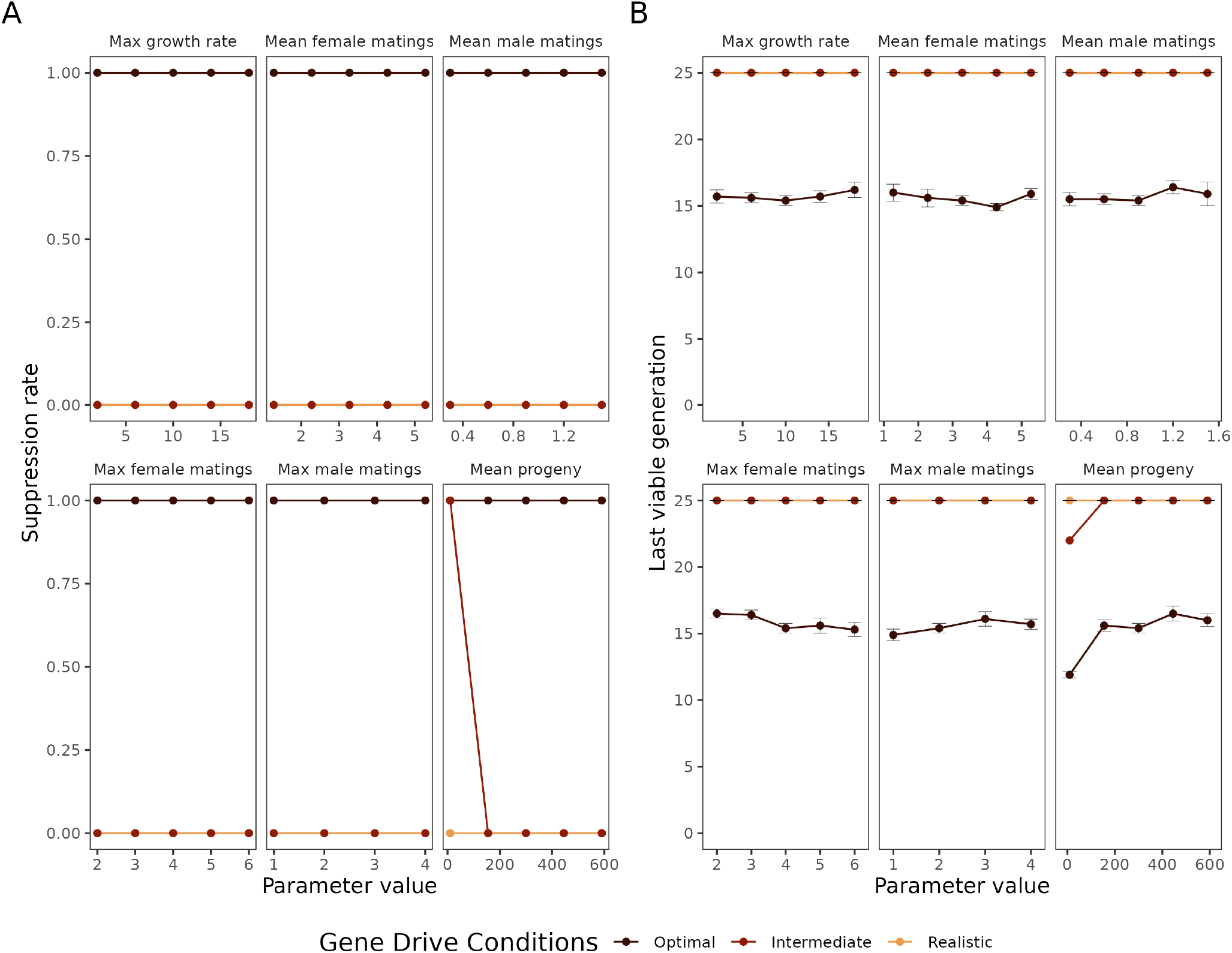
Sensitivity analysis for the biological parameters of the **Asian hornet**. **A** shows the suppression rate after the model has run for 25 generations. **B** shows the last viable generation over 25 years. The model was run under three different gene drive conditions: optimal, intermediate, and realistic. Under the realistic conditions the gene drive has a probability of non-homologous end-joining (*P_NHEJ_*) of 0.02, a cutting rate (*P_cut_*) of 0.95, and a heterozygous mortality (*P_mort_*) of 0.1. Under the optimal conditions these values are all 0. Under the intermediate conditions the gene drive has a probability of non-homologous end-joining (*P_NHEJ_*) of 0, a cutting rate (*P_cut_*) of 0.97, and a heterozygous mortality (*P_mort_*) of 0.15. Error bars represent the standard error of the mean.

**Fig. S4.**
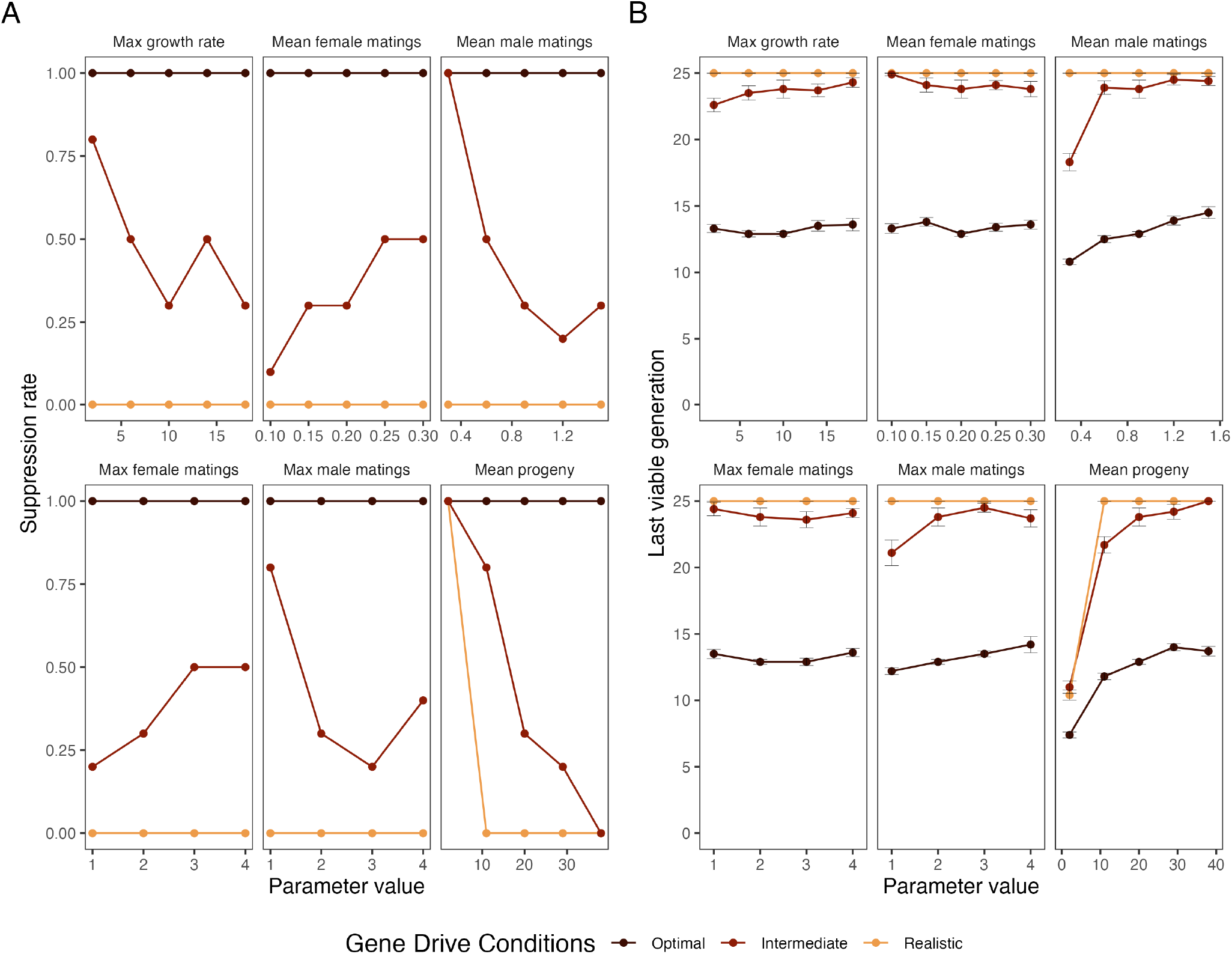
Sensitivity analysis for the biological parameters of the **European paper wasp A** shows the suppression rate after the model has run for 25 generations. **B** shows the last viable generation over 25 years. The model was run under three different gene drive conditions: optimal, intermediate, and realistic. Under the realistic conditions the gene drive has a probability of non-homologous end-joining (*P_NHEJ_*) of 0.02, a cutting rate (*P_cut_*) of 0.95, and a heterozygous mortality (*P_mort_*) of 0.1. Under the optimal conditions these values are all 0. Under the intermediate conditions the gene drive has a probability of non-homologous end-joining (*P_NHEJ_*) of 0, a cutting rate (*P_cut_*) of 0.97, and a heterozygous mortality (*P_mort_*) of 0.15. Error bars represent the standard error of the mean.

**Fig. S5.**
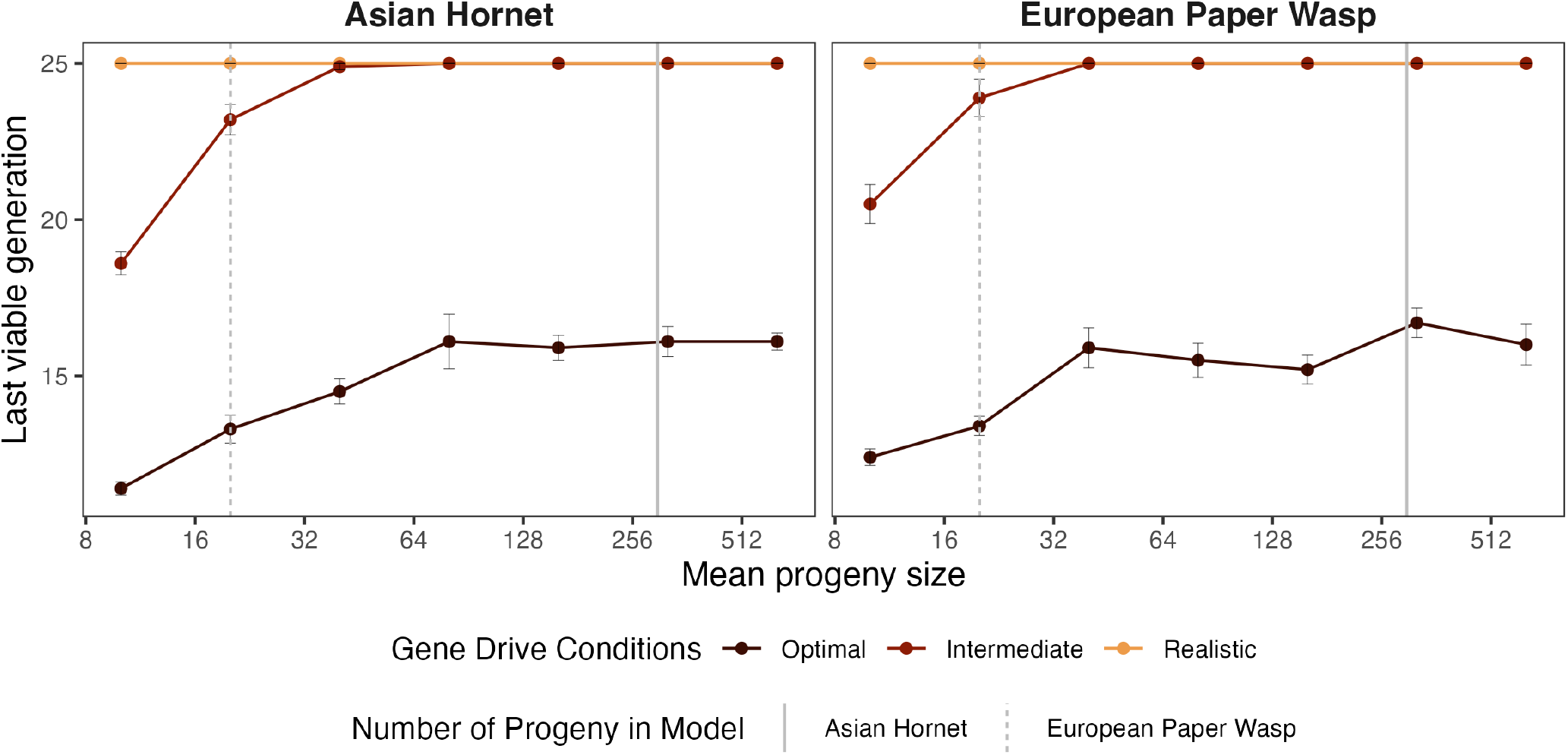
Last viable generation over 25 years for **A** The Asian hornet, and **B** the European paper wasp with different numbers of offspring. The grey lines indicate the values we used for each species. The model was run under three different gene drive conditions: optimal, intermediate, and realistic. Under the realistic conditions the gene drive has a probability of non-homologous end-joining (*P_NHEJ_*) of 0.02, a cutting rate (*P_cut_*) of 0.95, and a heterozygous mortality (*P_mort_*) of 0.1. Under the optimal conditions these values are all 0. Under the intermediate conditions the gene drive has a probability of non-homologous end-joining (*P_NHEJ_*) of 0, a cutting rate (*P_cut_*) of 0.97, and a heterozygous mortality (*P_mort_*) of 0.15. Error bars represent the standard error of the mean.

